# The outward Shaker channel OsK5.2 is beneficial to the plant salt tolerance through its role in K^+^ translocation and its control of leaf transpiration

**DOI:** 10.1101/2021.05.28.446164

**Authors:** Jing Zhou, Thanh Hao Nguyen, Doan Trung Luu, Hervé Sentenac, Anne-Aliénor Véry

**Affiliations:** BPMP, Univ Montpellier, CNRS, INRAE, Institut Agro, Montpellier, France

**Keywords:** Outward K^+^ channel, Shaker channel, salt tolerance, rice, K^+^/Na^+^ homeostasis, transpirational flux, xylem sap, root-to-shoot translocation, *Tos17* insertion mutants

## Abstract

High soil salinity constitutes a major environmental constraint to crop production worldwide, and the identification of genetic determinants of plant salt tolerance is awaited by breeders. While the leaf K^+^ to Na^+^ homeostasis is considered as key parameter of plant salt tolerance, the underlying mechanisms are not fully identified. Especially, the contribution of K^+^ channels to this homeostasis has been scarcely examined. Here, we show, using a reverse genetics approach, that the outwardly-rectifying K^+^ channel OsK5.2, involved in K^+^ translocation to the shoot and K^+^ release by guard cells for stomatal closure, is a strong determinant of rice salt tolerance. Upon saline treatment, OsK5.2 function in xylem sap K^+^ load was maintained, and even transiently increased, in roots. OsK5.2 selectively handled K^+^ in roots and was not involved in xylem sap Na^+^ load. In shoots, OsK5.2 expression was up-regulated from the onset of the saline treatment, enabling fast reduction of stomatal aperture, decreased transpirational water flow and therefore decreased trans-plant Na^+^ flux and reduced leaf Na^+^ accumulation. Thus, the OsK5.2 functions allowed shoot K^+^ nutrition while minimizing arrival of Na^+^, and appeared highly beneficial to the leaf K^+^ to Na^+^ homeostasis, the avoidance of salt toxicity and plant growth maintaining.

## 1 INTRODUCTION

High soil salinity is a widespread environmental constraint over the world that causes substantial restrictions in production and quality of a majority of crops, including cereals. Understanding how plants cope with high salinity in the environment is thus an issue of great agricultural importance. Rice is rated as a salt-sensitive cereal (Munns & Tester, 2008; Zeng & Shannon, 2000), and salinity levels have increased in rice fields, in particular owing to the climate changes and sea level rise, which strongly challenge rice culture in coastal regions.

The adverse effects of high soil salinity on plant growth are mainly related to the decrease in osmotic potential of the soil and to ionic toxicity of Na^+^ in leaves (Munns & Tester, 2008). Studies on the latter phenomenon have revealed that genes encoding Na^+^ transport systems correspond to major quantitative trait loci (QTLs) of salt tolerance (Hauser & Horie, 2010). In return, such findings have spurred research efforts in this domain of membrane transport biology, highlighting also that Na^+^ detrimental effects are counteracted by the plant’s ability to take up the essential macronutrient K^+^ and control its K^+^ nutritional status in presence of high external Na^+^ concentrations (Maathuis & Amtmann, 1999). K^+^ is involved in vital functions such as enzyme activation, the cytoplasmic pH homeostasis, control of cell membrane potential and cell turgor-driven movements (Marschner, 2011; Nieves-Cordones, Al Shiblawi & Sentenac, 2016). Upon salt stress, the massive influx of positively charged Na^+^ ions causes cell membrane depolarization, which reduces the driving force for K^+^ uptake and even in some cases leads to channel-mediated root K^+^ losses (Jayakannan, Bose, Babourina, Rengel, & Shabala, 2013; Rubio, Nieves □ Cordones, Horie, & Shabala, 2020). Thus, plant exposure to high salinity is inevitably accompanied by chronic K^+^ deficiency, which affects the leaf K^+^ to Na^+^ content ratio, whose maintenance to a high value is a key determinant of salt tolerance (Hauser & Horie, 2010; Maathuis & Amtmann, 1999).

K^+^ and Na^+^ ions taken up by root cells can migrate to stelar tissues and be translocated to leaves by the upward flow of sap in the xylem vessels. Control of the ionic composition of xylem sap, involving membrane ion transport processes in parenchyma cells along the sap ascent pathway, is thus a major determinant of salt tolerance, together with control of the flux of xylem sap, which is driven by leaf transpiration and hence dependent on the level of stomatal aperture, or driven by the so-called root pressure, resulting from increased osmotic pressure in the xylem vessels due to increased solute concentration in the sap in absence of significant plant transpiration (Jeschke, 1984; Marschner, 2011). Therefore, transport systems contributing to Na^+^ or K^+^ secretion/retrieval into/from the xylem sap or to regulation of stomatal aperture can contribute to processes that play crucial roles in salt tolerance.

In various plant species, Na^+^ transporters from the HKT family have been shown to contribute to Na^+^ retrieval from the xylem sap and loading into the xylem parenchyma cells bordering the vessels, *i.e.* to the so-called “sap desalinization” process (Hauser & Horie, 2010). In rice, the *HKT* transporter genes identified as involved in this process are *OsHKT1;5,* which is mainly expressed in root xylem parenchyma cells and corresponds to the major salt-tolerance QTL *SKC1* (Ren et al., 2005), *OsHKT1;4,* expressed in both root and basal leaf xylem tissues (Suzuki et al., 2016; Khan et al., 2020), and *OsHKT1;1* expressed in both xylem and phloem and thereby additionally involved in Na^+^ recirculation from leaves to roots within the phloem sap, favoring root *versus* leaf Na^+^ accumulation (Campbell et al., 2017; Wang et al., 2015).

Compared with the large number of studies focused on the Na^+^ transporters controlling Na^+^ translocation to leaves and thereby contributing to maintain the ratio of leaf K^+^ to Na^+^ contents to a high value, less attention has been paid to the K^+^ transport mechanisms that operate under saline conditions and ensure efficient K^+^ supply to leaves. Current knowledge in this domain essentially concerns K^+^ transport systems involved in root K^+^ uptake, and mainly high-affinity K^+^ transporters from the HAK/KUP/KT family, AtHAK5 in Arabidopsis and OsHAK1, OsHAK5, OsHAK16 and OsHAK21 in rice (Chen et al., 2015; Feng et al., 2019; Nieves-Cordones, Alemán, Martínez & Rubio, 2010; Shen et al., 2015; Yang et al., 2014).

In rice, the outwardly rectifying Shaker K^+^ channel OsK5.2 is involved both in K^+^ translocation into the xylem sap toward the shoots and in control of stomatal aperture and leaf transpiration by driving K^+^ efflux from guard cells for stomatal closure (Nguyen et al., 2017). This channel thus emerged as a good model to assess the level of contribution of these functions to the control of Na^+^ and K^+^ delivery to shoots upon saline conditions and salt tolerance. This has been achieved in the present study by phenotyping *osk5.2* knock-out (KO) mutant plants subjected to saline conditions. We found that the lack of functional *OsK5.2* expression does result in increased plant sensitivity to salt stress and analyzed the bases of the salt sensitive phenotype.

## 2 MATERIALS AND METHODS

### 2.1 Plant growth and salt treatment

The selection from *Tos17*-insertion lines of *osk5.2* mutant and corresponding wild-type (WT) plants in the background of rice Nipponbare cultivar *(Oryza sativa* L. ssp. japonica cv. Nipponbare) has been previously described (Nguyen et al., 2017). Rice seeds were germinated on a raft floating on deionized water for one week. The seedlings were then hydroponically grown on half-strength Yoshida medium for one week, and thereafter on Yoshida medium (0.5 mM (NH_4_)_2_SO_4_, 1.6 mM MgSO_4_, 1.2 mM Ca(NO_3_)_2_, 0.7 mM KNO_3_, 0.8 mM KH_2_PO_4_, 60 μM Na_2_FeEDTA, 20 μM MnSO_4_, 0.32 μM (NH_4_)_6_Mo_7_O_24_, 1.4 μM ZnSO_4_, 1.6 μM CuSO_4_, 45.2 μM H_3_BO_3_, and pH adjusted to 5.5 with H_2_SO_4_). Five-week-old rice plants were subjected to salt treatment by supplementing the hydroponic Yoshida medium with NaCl. The rice plants were grown in a growth chamber (70% relative humidity, light intensity 130 photon μmol.m^-2^.s^-1^, 29°C/25°C 12 h/12 h day/night).

### 2.2 RNA extraction and quantitative real time PCR experiments

Five-week-old Nipponbare plants grown on Yoshida medium were either supplemented with 50 mM NaCl for 14 days and then transferred back to standard Yoshida medium for three days, or further grown during this period on Yoshida medium (control batch). Total RNAs were extracted from samples collected at same times from salt-treated or control plant batches using the RNeasy plus mini kit with gDNA eliminator (Qiagen, Germany). First-strand cDNAs were synthesized from 3 μg of RNAs using SuperScript III reverse transcriptase (Invitrogen) and used as template for qRT-PCR experiments. qRT-PCR analyses were performed using the Lightcycler480 system (Roche diagnostics) and SYBR *Premix Ex Taq* (Takara) in a total volume of 10 μl, which contained 2 μl of cDNA, 3 μl of forward and reverse primer mixture (1 μM), and 5 μl of SYBR *Premix Ex Taq.* Reactions were performed with three independent biological replicates, each one with three technical replicates (PCR program: 95°C for 30 sec; 45 cycles of 95°C for 10 sec, 60°C for 10 sec, and 72°C for 15 sec; followed by a melt cycle from 60°C to 95°C). C_T_ (cycle threshold) values were obtained from amplification data using a threshold of 0.37. The *OsK5.2* absolute number of copies was calculated according to standard curves obtained by successive dilutions with known quantities of *OsK5.2,* and then normalization using the C_T_ values of three housekeeping genes (ubiquitin-like protein gene *SMT3,* PP2A-interactor gene *Tip41* and elongation factor gene *EF1 beta*) as described in Khan et al. (2020). The sequences of the primers used for qRT-PCR experiments are provided in Table S1.

### 2.3 Na^+^ and K^+^ assays in tissues and xylem sap

Five-week-old plants hydroponically grown as described above were supplemented or not with 50 mM NaCl for 14 days. Excised root systems and shoots were periodically collected during the salt treatment, both from salt treated and control plants. The roots (rinsed in deionized water) and shoots were dried (60 □ for 3 days) and weighed. Ions were extracted from the tissues in 0.1 N HCl for 3 days and assayed by flame spectrophotometry (SpectrAA 220FS, Varian).

Xylem sap was collected through natural exudation under control condition from de-topped plants (3 cm above the root system). Upon salt treatment, xylem sap was obtained through pressurization. The root system of the de-topped plants was placed into a pressure chamber (Boursiac *et al.,* 2005) filled with hydroponic medium containing 50 mM NaCl, and sealed with a silicon dental paste (PRESIDENT Light Body, Coltene, Switzerland). The first few drops (2 μl) were discarded to avoid the contamination that results from injured cells. Twenty μl of sap samples were then collected using a micro-pipette, transferred into 0.2 ml Eppendorf tubes kept on ice, and diluted in 0.1 N HCl for Na^+^ and K^+^ assay (flame spectrophotometry).

### 2.4 Leaf transpiration

Intact plants were transferred into a multipotometer device (Nguyen et al., 2017) one day before transpiration rate measurement. The root system of each intact plant was inserted into a 50 ml syringe filled with hydroponic medium through the rubber plunger, and sealed with a silicon dental paste. Each syringe was connected to a graduated 1-ml serological plastic pipette via a thin silicone tube. The pipettes were refilled with the same medium with proper time intervals, making sure that no air bubble was present in the system. A camera took photographs of the set of pipettes every two minutes in order to record the changes in water level in the pipettes. Image capture was started after 3 h under light condition and maintained for 30 minutes. Plants were then exposed to darkness for 5 h under continuous recording. The rate of decrease in the water volume in the pipettes was used to calculate the mean transpiration rate of plants.

## 3 RESULTS

### 3.1 *osk5.2* mutant plants exhibit increased sensitivity to salinity

The two *osk5.2* KO mutant lines ASJA08 and ASHF06 (Nguyen et al., 2017) and the corresponding wild type (WT) plants displayed a very similar development and phenotype when growth occurred in absence of salt stress (Figure 1). This was no longer the case when the plants were subjected to saline conditions (6-week-old plants subjected to 100 mM NaCl for 7 days). The mutant plants then displayed a severe reduction in biomass, by about 45-50% when compared with the biomass of plants from the same mutant lines but grown in control conditions, while the corresponding reduction observed for the WT plants appeared weak (less than 10-15%) and not statistically significant (Figure 1b). Furthermore, under this saline treatment, the mutant plants displayed a greater extent of stunted leaf growth and a larger number of dried leaves than the WT plants (Figure 1a). Thus, the lack of *OsK5.2* functional expression resulted in reduced tolerance to saline conditions.

**FIGURE 1.**
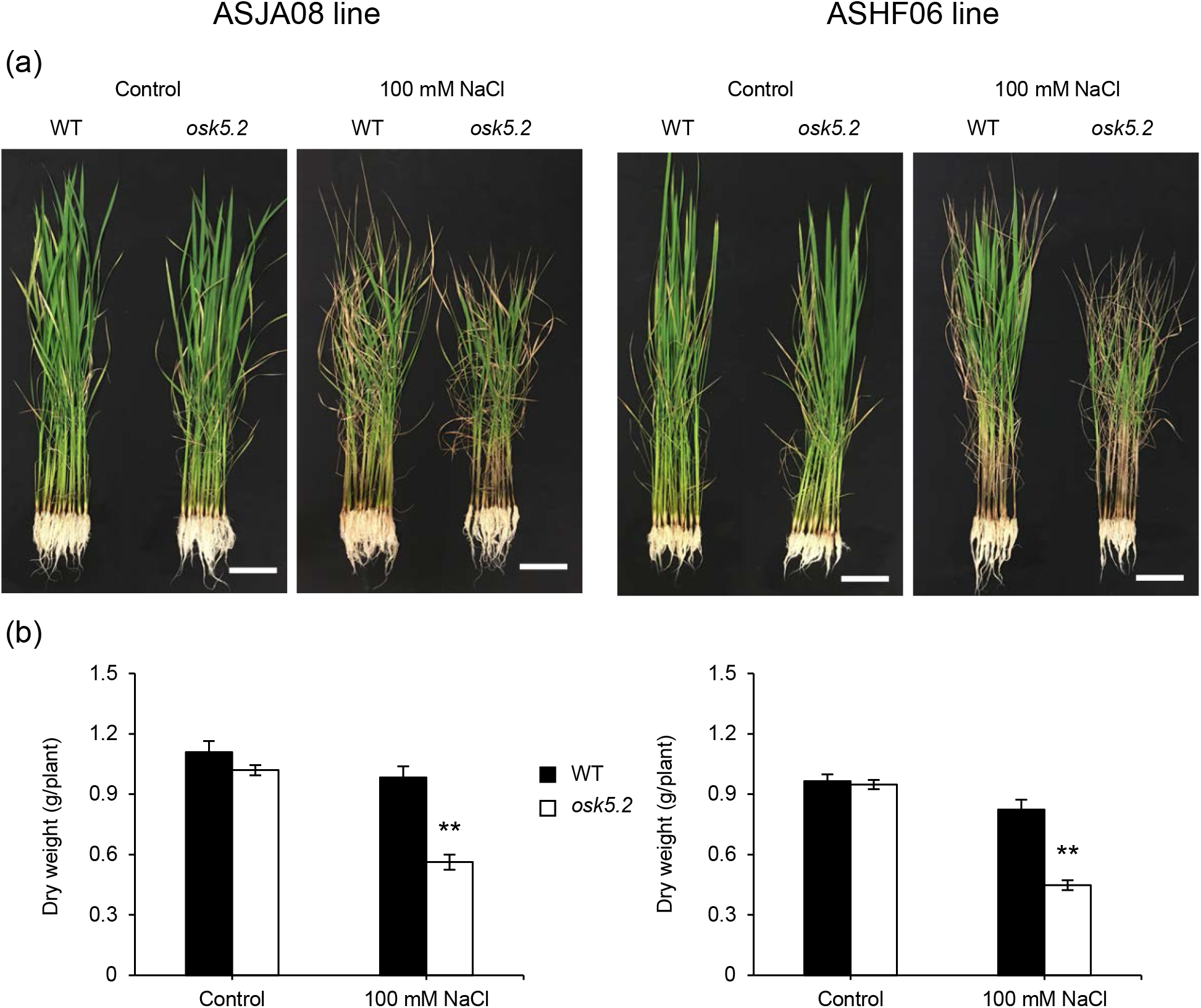
Effect of *OsK5.2* loss of function on rice plant phenotype in control and salt stress conditions. Comparison of growth phenotype (a) and dry weight (b) between corresponding wild-type and *osk5.2* mutant plants (black and white bars, respectively) issued from ASJA08 or ASHF06 lines (left and right panels, respectively) under control and salt treatment. Six-week-old plants grown on hydroponic Yoshida medium were supplemented or not during the last 7 days with 100 mM NaCl. Scale bars = 10 cm in (a). Means ± SE, *n* = 10. Double stars above the bars denote statistically significant differences between wild-type and *osk5.2* mutant plants (Student’s *t* test, *P* ≤ 0.01).

### 3.2 *OsK5.2* transcript accumulation under salt stress

Real-time qRT-PCR analyses revealed that *OsK5.2* was expressed in both roots and leaves (Figure 2). In plants grown in control condition, *OsK5.2* transcripts were 4 folds more abundant in roots than in leaves. When the plants were subjected to saline conditions (50 mM NaCl), a change in the relative expression of *OsK5.2* between roots and shoots was observed leading to balanced expression in the plant. In leaves, the accumulation of *OsK5.2* transcripts was rapidly (from one day after salt treatment) up-regulated by about 3 folds, and remained high during 7 days (Figure 2b). The leaf level of *OsK5.2* transcripts appeared to decrease with longer exposure to the saline conditions but, after 14 days of salt treatment, it was still about 1.5-fold that observed in control conditions (Figure 2b). Recovery from salt stress for 1 to 3 days further decreased the leaf level of *OsK5.2* transcripts, down to that observed in leaves from control plants. In roots, as compared with leaves, the accumulation of *OsK5.2* transcripts showed more reduced variations in response to the saline treatment and tended to slightly decrease, a down-regulation by about 25% observed at days 3 and 14 of the salt treatment being statistically significant (Figure 2a). After 1 day of recovery from salt stress, the root level of *OsK5.2* transcripts recovered to same expression level of control plants and then remained stable for at least 2 days.

**FIGURE 2.**
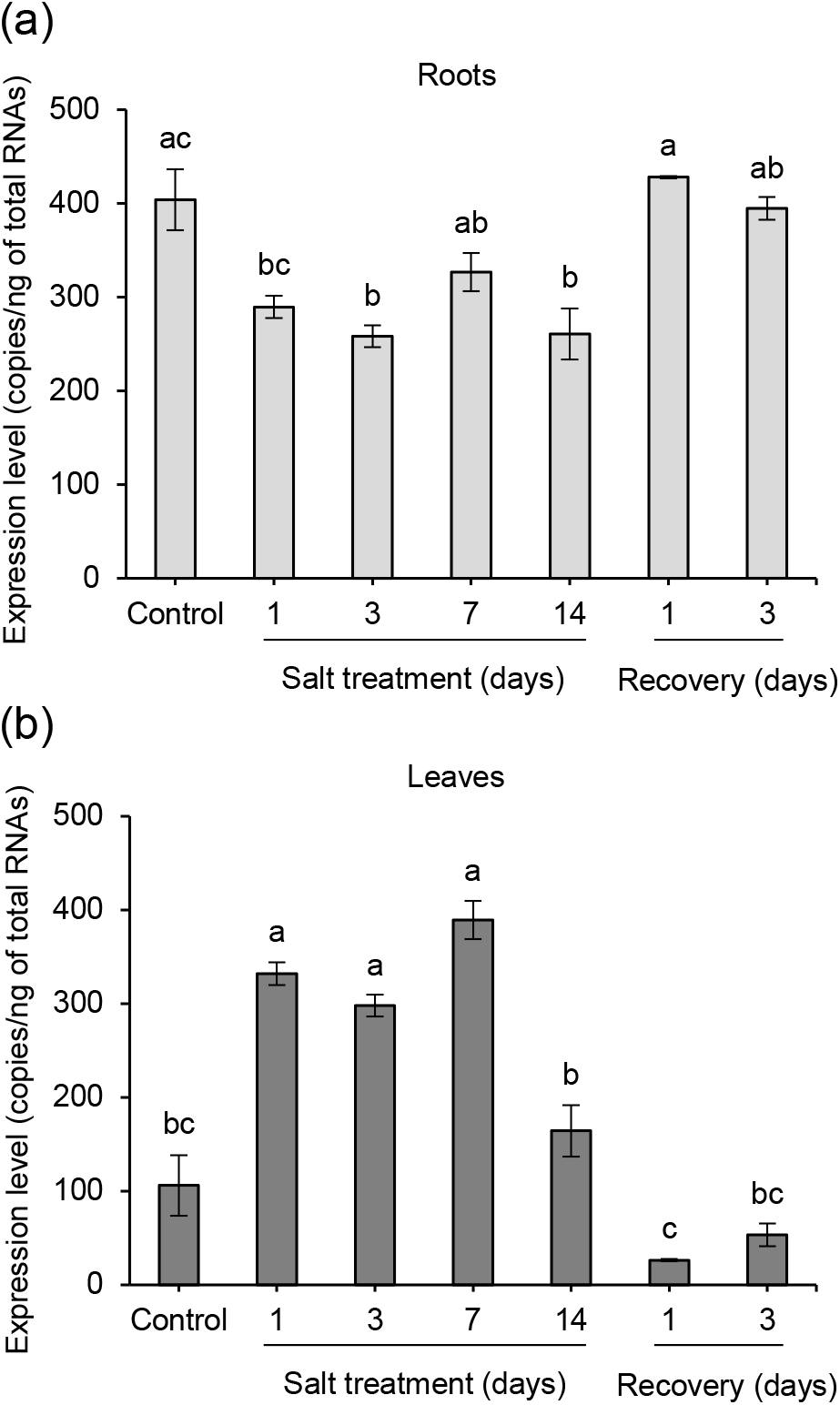
Effect of saline conditions on *OsK5.2* transcript levels in roots and leaves. Five-week-old rice plants cv Nipponbare hydroponically grown on Yoshida medium were supplemented or not with 50 mM NaCl for 14 days. Salt-treated plants were thereafter allowed to recover for 3 days on standard Yoshida medium. Expression data in roots (a) and leaves (b) were determined by real-time quantitative RT-PCR. Means ± SE (*n* = 3 biological replicates under salt treatment after 1, 3, 7 and 14 days and recovery, and *n* = 4 under control treatment sampled at each time of salt treatment). Different letters indicate statistically significant differences (Student’s *t* test, *P* ≤ 0.05).

### 3.3 *osk5.2* mutant plants display larger transpirational water loss than WT plants under salt stress

To investigate the role of *OsK5.2* in control of leaf transpiration under salt stress, 50 mM NaCl was added into the hydroponics solution of 5-week-old *osk5.2* mutant and WT plants, and the steady-state rates of plant water loss were measured periodically for 14 days (on days 1, 3, 7 and 14 of the salt treatment) both under light and in dark conditions using a multipotometer (Figure 3). The data obtained with the 2 *osk5.2* mutant lines ASJA08 and ASHF06 led to the same conclusions. The salt treatment strongly decreased the rate of transpirational water loss, in the *osk5.2* mutant and the corresponding WT plants, by up to 50 to 55 % in the light and 30 to 40% in the dark conditions (Figure 3). The kinetics of reduction of transpiration rate upon NaCl exposure was clearly slower in *osk5.2* mutant compared with WT plants both in light and dark conditions. During the two weeks of salt treatment, a higher rate of transpirational water loss was consistently observed in *osk5.2* mutant as compared with WT plants in light and dark conditions, and the difference was highly significant in most of the analyzed time points (Figure 3). The greatest difference in transpiration rate between WT and *osk5.2* mutant plants occurred after one day of treatment owing to the slower response to NaCl exposure in *osk5.2* mutant plants.

**FIGURE 3.**
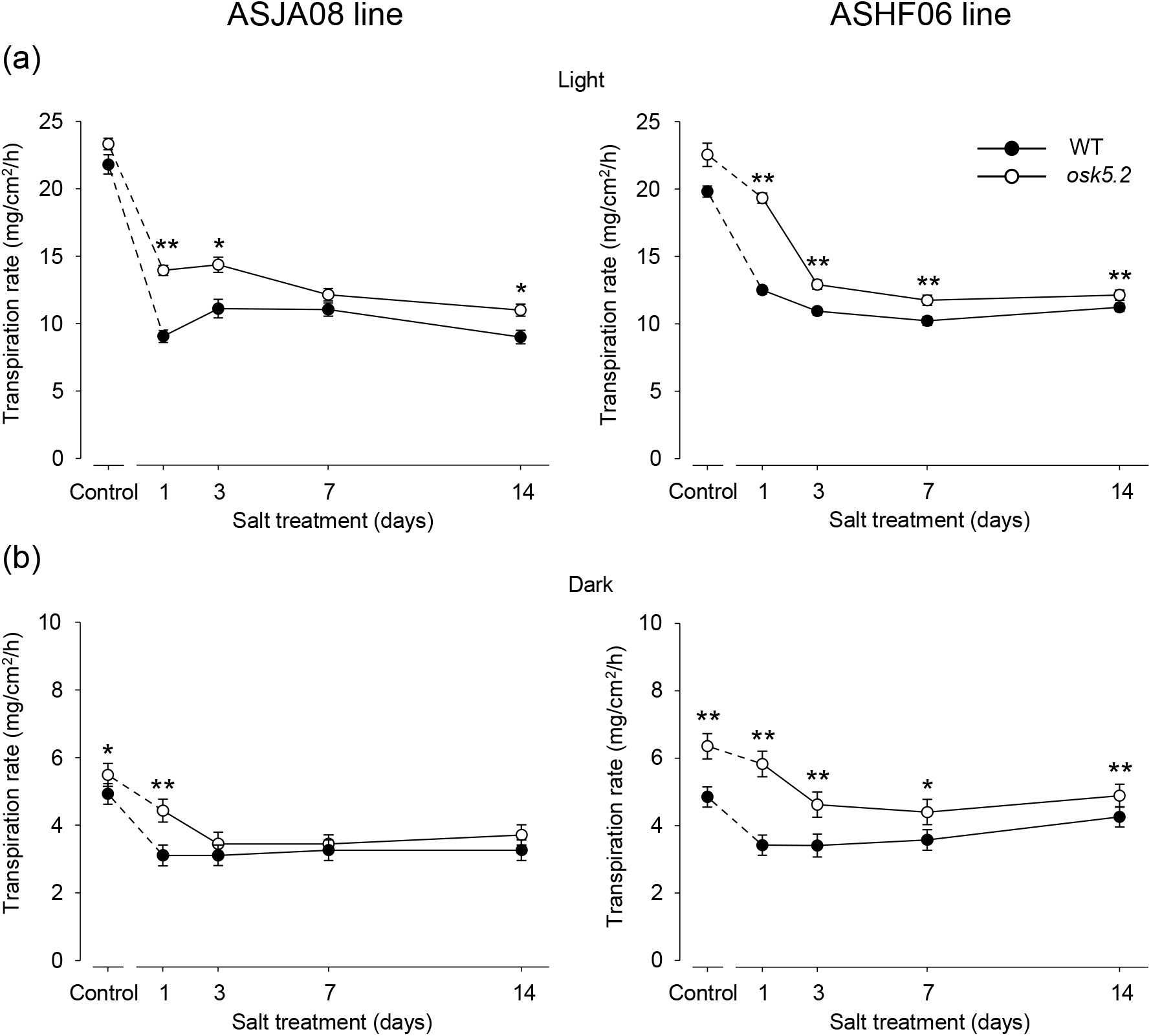
Steady-state transpiration rates in wild-type and *osk5.2* mutant plants under control and salt treatment conditions. Five-week-old plants hydroponically grown on Yoshida medium were supplemented or not with 50 mM NaCl for 14 days. Left and right panels: *osk5.2* mutant plants (◦) issued from ASJA08 or ASHF06 lines, respectively, and the corresponding wild-type plants (•). Transpiration was measured after 1, 3, 7 and 14 days of salt treatment (and at the same times for the plants maintained in control conditions). (a) and (b): steady-state transpiration rates in light (panel a; ~3 h after light was switched on) and in dark (panel b; ~5 h after light was switched off) conditions. Steady-state transpiration rate was determined by dividing the average plant rate of water loss at steady-state (means of 3 values) by the total surface of the plant aerial parts. Means ± SE; *n* = 9 under salt treatment after 1, 3, 7, 14 days, and *n* = 12 under control conditions. Single and double stars denote statistically significant differences between wild-type and *osk5.2* mutant plants (Student’s *t* test, *P* ≤ 0.05 and *P* ≤ 0.01, respectively).

### 3.4 Na^+^ and K^+^ concentrations in xylem sap under salt stress and translocation fluxes towards the shoots

Xylem sap samples were collected from de-topped plants subjected to the same protocol of salt treatment as that that used in the experiment described by Figure 3: 2 weeks in 50 mM NaCl hydroponics solution applied to 5-week-old plants previously grown under control conditions. The concentrations of K^+^ and Na^+^ determined in sap samples (Figure 4) and the transpiration rates recorded under the same experimental conditions (Figure 3) were used to estimate the transpiration-driven fluxes of K^+^ and Na^+^ (Figure S1) arriving in shoots under such conditions.

**FIGURE 4.**
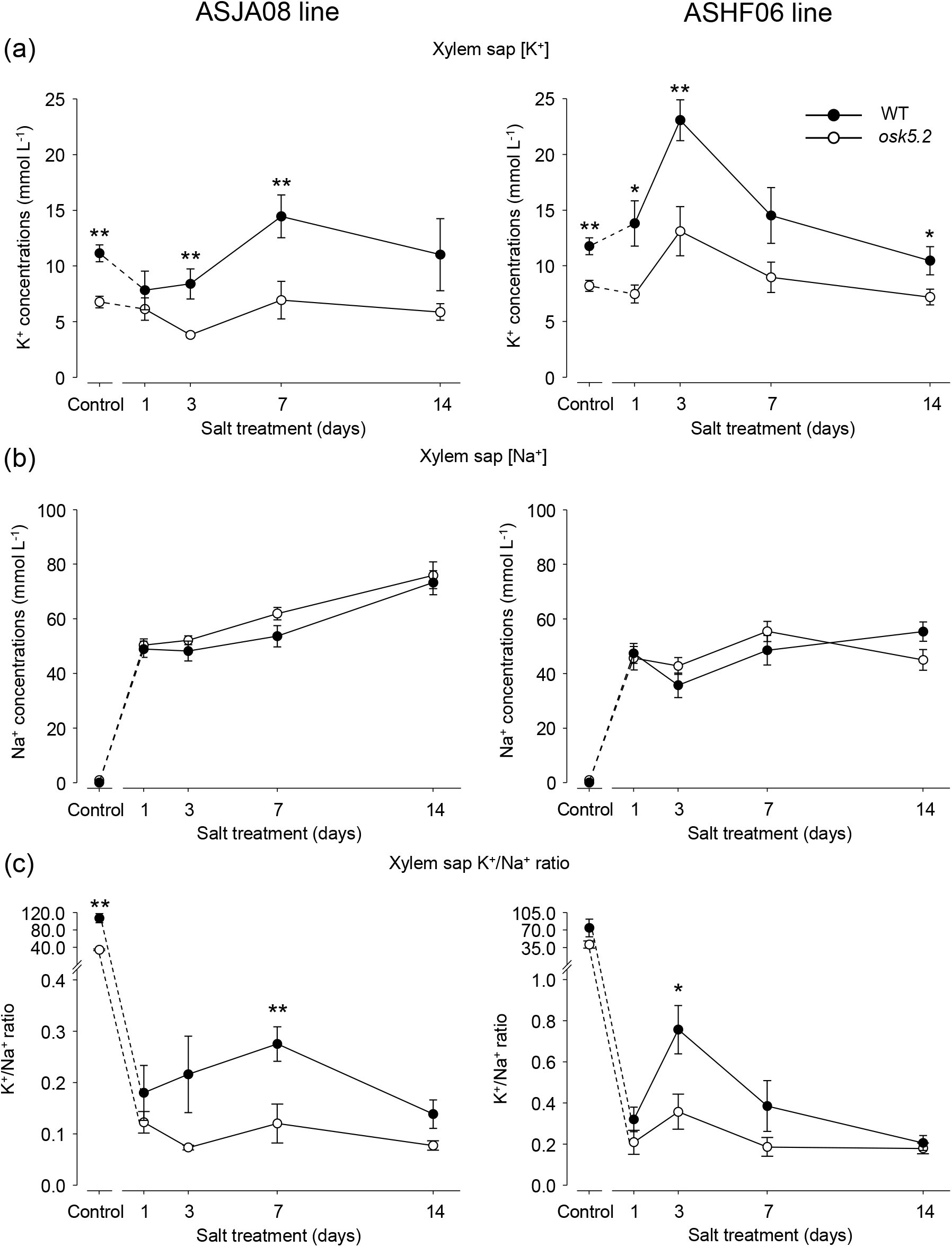
Xylem sap K^+^ and Na^+^ concentrations in wild-type and *osk5.2* mutant plants under control and salt treatment conditions. Five-week-old plants hydroponically grown on Yoshida medium were supplemented or not with 50 mM NaCl for 14 days. Left and right panels: *osk5.2* mutant plants (◦) issued from ASJA08 or ASHF06 lines, respectively, and the corresponding wild-type plants (•). Xylem sap exudates were collected after 1, 3, 7 and 14 days of salt treatment (and at the same times for the plants maintained in control conditions). (a) and (b): K^+^ (a) and Na^+^ (b) concentrations assayed in the collected xylem sap samples. (c) K^+^/Na^+^ concentration ratios deduced from (a) and (b). Means ± SE; *n* = 9 under salt treatment after 1, 3, 7, 14 days, and *n* = 12 under control conditions. Single and double stars denote statistically significant differences between wild-type and *osk5.2* mutant plants (Student’s *t* test, *P* ≤ 0.05 and *P* ≤ 0.01, respectively).

Under control condition, K^+^ concentrations measured in xylem sap were close to 11 mM (7-folds the K^+^ concentration in the hydroponic medium) in WT plants, and were 30 to 40% lower in the two *osk5.2* mutant lines (as previously reported; Nguyen et al., 2017). The xylem sap K^+^ concentration in both WT and *osk5.2* mutant plants displayed transient variations upon exposure to the saline conditions (Figure 4a). The concentrations observed in the *osk5.2* mutant plants remained lower than those displayed by the corresponding WT plants by about 40% to 50% over the entire duration of the salt treatment (Figure 4a). With respect to Na^+^, the concentration of this cation in the xylem sap was extremely low, in the submillimolar range, in the absence of salt treatment (Figure 4b). Exposure to 50 mM NaCl led to the loading of a large amount of Na^+^ to the xylem sap with no significant difference between WT and *osk5.2* mutant plants (Figure 4b). The Na^+^ concentrations measured in the xylem sap of the two types of plants were close to that of the hydroponic medium (50 mM) after one day of NaCl supplementation, and remained fairly stable during the two weeks of salt treatment. Altogether, these results indicated that the lack of *OsK5.2* functional expression constitutively resulted in a large reduction in xylem sap K^+^ concentration, by *ca.* 40-50%, but did not affect xylem sap Na^+^ concentration. As a result, the K^+^/Na^+^ xylem sap concentration ratios, computed from the data provided by Figure 4a and 4b, appear consistently higher in WT than in *osk5.2* mutant plants (Figure 4c).

The estimated K^+^ flux arriving in shoots during the light period in these experimental conditions, obtained by integrating the data from Figure 3a and 4a (Figure S1a), is decreased by the salt treatment in both the *osk5.2* mutant and WT plants. It is lower in both *osk5.2* mutant lines than in the corresponding WT plants, except at day 1 of the salt treatment due to the sharp decrease in transpiration rate displayed by WT plants at this time point. The differences between the WT and mutant plants under salt treatment are in the 30 to 50% range from days 3 of the salt treatment, *i.e.,* similar to those observed between the two types of plants in control conditions (Figure S1a).

The estimated flux of Na^+^ arriving in the shoots of WT and *osk5.2* mutant plants is very low under control conditions (this cation being then present as trace contaminant in the hydroponics solution). A marked increase in Na^+^ flux is observed in all genotypes from the first day of exposure to NaCl (Figure S1b). The flux is larger in *osk5.2* mutant than in WT plants, by ca. 20 to 40% during the first week of the salt treatment, which essentially reflects the difference in transpiration rate between the two types of plants during this period (see Figure 3).

### 3.5 *osk5.2* mutant plants accumulate less K^+^ and more Na^+^ under salt stress

K^+^ and Na^+^ contents were determined in roots and shoots of *osk5.2* and WT plants subjected to the same salt treatment protocol as that previously used.

In all genotypes, root and shoot K^+^ contents, and thus whole plant K^+^ contents, decreased with the duration of the salt treatment (Figure 5a, b, respectively). In shoots, the decrease was clearly more pronounced in the *osk5.2* mutant lines, when compared with the corresponding WT plants, and the relative difference in shoot K^+^ contents between the mutant and WT plants increased with the duration of the salt treatment (Figure 5b): the difference was in the range of 10-20% at the beginning of the treatment (in the absence of NaCl addition and at day 1 of the salt treatment), and reached 40-60% after two weeks of treatment (Figure 5b). In roots, the impact of lack of *OsK5.2* functional expression on K^+^ contents appeared much weaker than that observed in shoots (Figure 5a).

**FIGURE 5.**
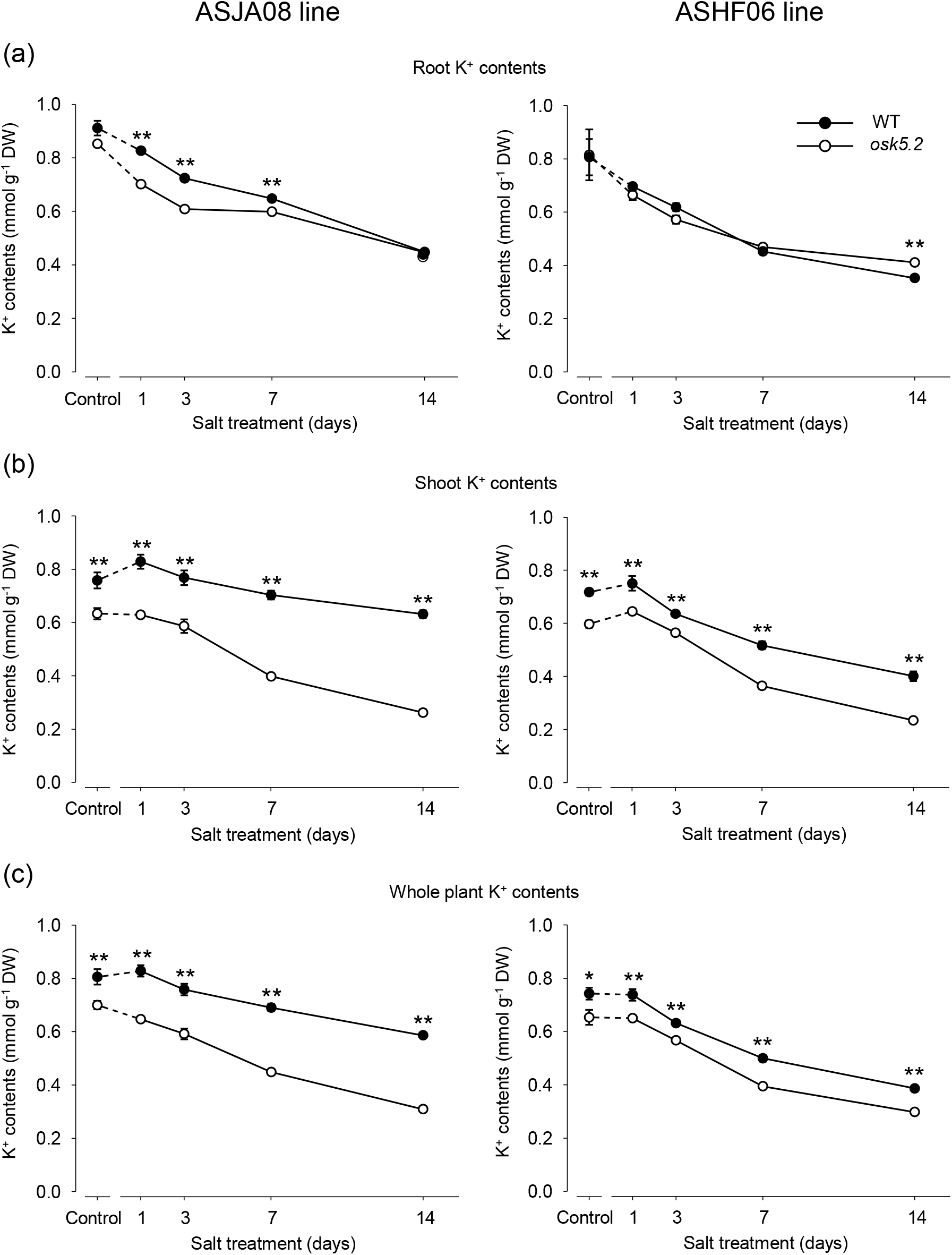
Root and Shoot K^+^ contents in wild-type and *osk5.2* mutant plants under control and salt treatment conditions. Five-week-old plants hydroponically grown on Yoshida medium were supplemented or not with 50 mM NaCl for 14 days. Left and right panels: *osk5.2* mutant plants (◦) and the corresponding wild-type plants (•) issued from ASJA08 (left) or ASHF06 (right) lines. Roots and shoots were sampled after 1, 3, 7 and 14 days of salt treatment (and at the same times for the plants maintained in control conditions). (a), (b) and (c): K^+^ contents in roots, shoots and whole plant, respectively. Means ± SE; *n* = 9 under salt treatment after 1, 3, 7 and 14 days, and *n* = 12 under control conditions. Single and double stars denote statistically significant differences between the wild-type and *osk5.2* mutant plants (Student’s *t* test, *P* ≤ 0.05 and *P* ≤ 0.01, respectively).

Regarding Na^+^, the contents of this cation were very low in all plants, whatever their genotype, in the absence of salt treatment (Figure 6). The salt treatment increased both the root and shoot (and thus the whole plant) contents of this cation, from the first day of treatment and over the two weeks of treatment, in all plant genotypes (Figure 6). Under all conditions except the longest duration of the salt treatment (in other words, under control conditions and during the first week of salt exposure), significantly higher Na^+^ contents were found in the *osk5.2* mutants than in the corresponding WT plants, for both mutant lines, by more than 20% in roots and 35% in shoots (Figure 6b). The relative differences between the mutant and WT plants were weaker at the last time point of salt treatment (after 14 days), and the differences were no longer statistically significant (except for the roots of one mutant line). At this time, Na^+^ levels in shoots exceeded those in roots in all genotypes (Figure 6a, b). Thus, Na^+^ accumulation was higher in *osk5.2* mutant plants than in the corresponding WT plants until the late stage of Na^+^ plant invasion, when the level of Na^+^ in shoots had become higher than that in roots.

**FIGURE 6.**
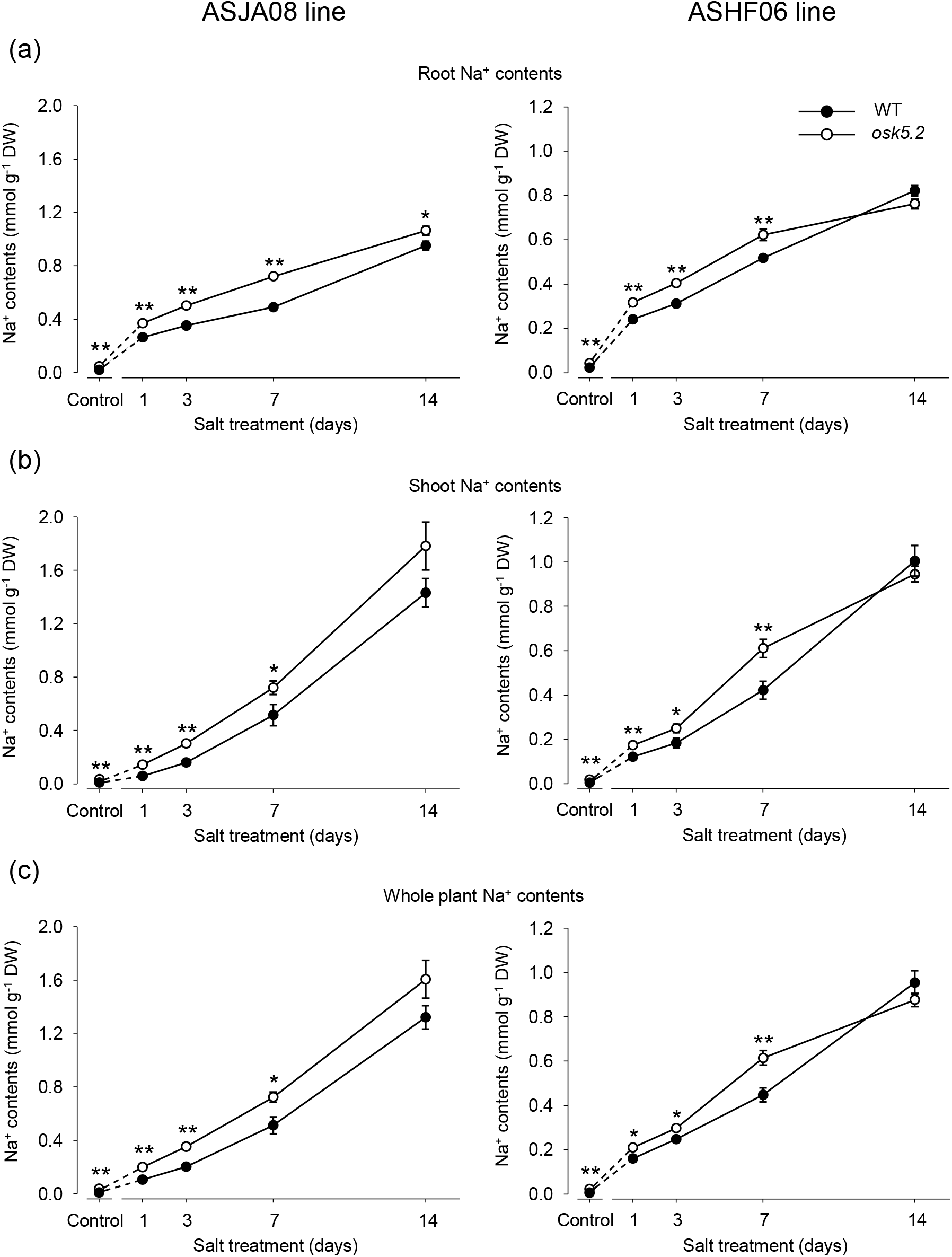
Root and Shoot Na^+^ contents in wild-type and *osk5.2* mutant plants under control and salt treatment conditions. Same plants as in Figure 5. (a), (b) and (c): Na^+^ contents in roots, shoots and whole plant, respectively. Means ± SE; *n* = 9 under salt treatment after 1, 3, 7 and 14 days, and *n* = 12 under control conditions. Single and double stars denote statistically significant differences between the wild-type and *osk5.2* mutant plants (Student’s *t* test, *P* ≤ 0.05 and *P* ≤ 0.01, respectively).

K^+^/Na^+^ content ratios were calculated from the data displayed by Figures 5 and 6. The ratios were significantly lower in *osk5.2* mutant plants than in WT plants both in shoots and roots under all conditions except in roots of one mutant line at the last time point of salt treatment (Figure 7). In salt-stressed leaves, the relative reduction in K^+^/Na^+^ content ratios observed in the *osk5.2* mutant plants as compared with the corresponding WT plants, was in the range of 35-70% (Figure 7). Thus, *OsK5.2* lack of functional expression strongly impaired K^+^/Na^+^ homeostasis in leaves under salt stress.

**FIGURE 7.**
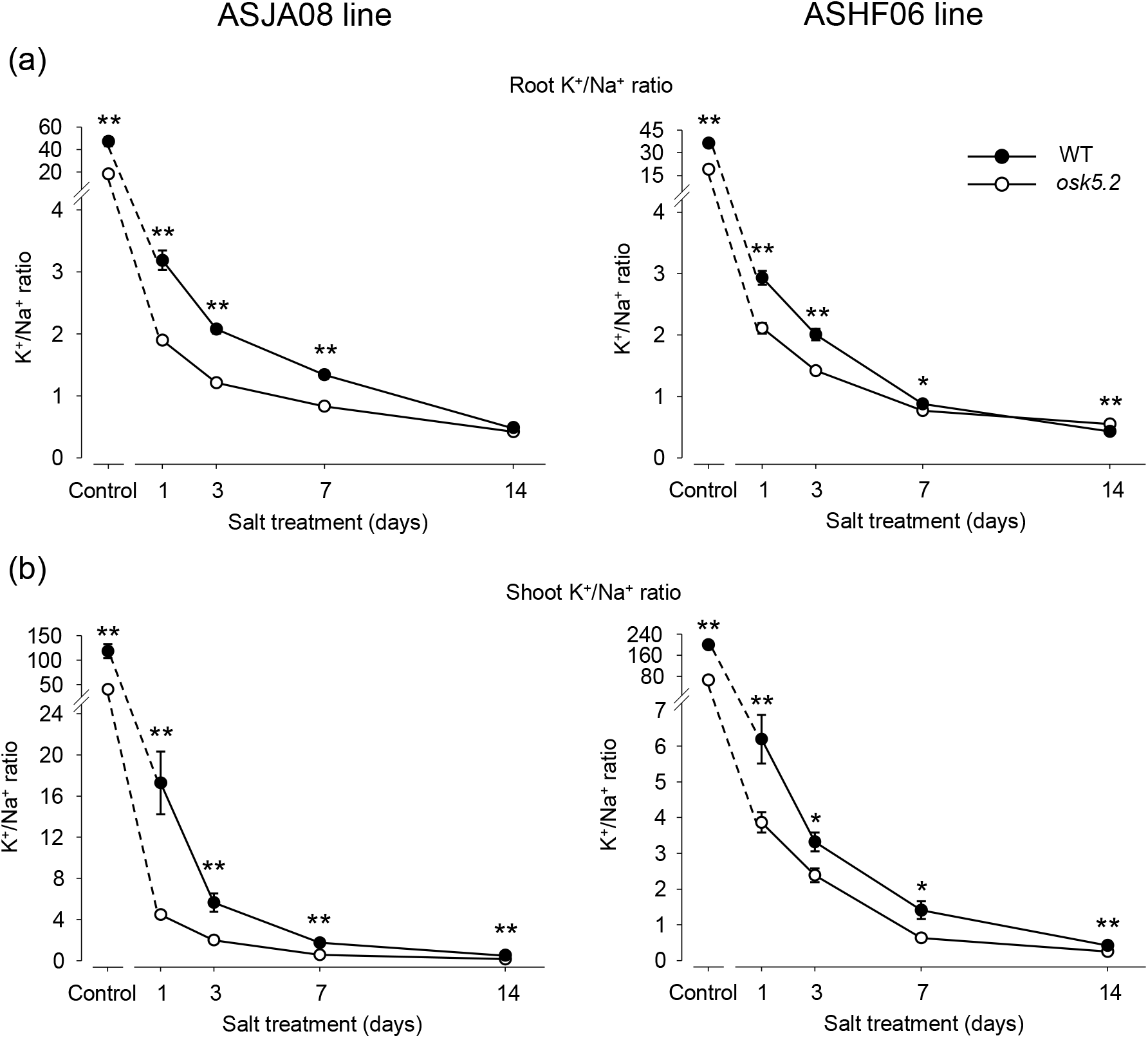
Root and shoot K^+^/Na^+^ content ratio in wild-type and *osk5.2* mutant plants under control and salt treatment conditions. Same experiment as in Figures 5 and 6. K^+^/Na^+^ content ratio: K^+^ content from Figure 5 divided by the corresponding Na^+^ content from Figure 6. (a) and (b): K^+^/Na^+^ content ratio in roots and shoots. Left and right panels: *osk5.2* mutant plants (◦) and corresponding wild-type plants (•) issued from ASJA08 (left) or ASHF06 (right) lines. Means ± SE; *n* = 9 under salt treatment after 1, 3, 7 and 14 days, and *n* = 12 under control conditions. Single and double stars denote statistically significant differences between the wild-type and *osk5.2* mutant plants (Student’s *t* test, *P* ≤ 0.05 and *P* ≤ 0.01, respectively).

## 4 DISCUSSION

### 4.1 OsK5.2, a model for analyzing K^+^ channel-mediated control of K^+^ and Na^+^ translocation to the shoots under salt stress

Mechanisms that control long distance transport of Na^+^ and K^+^ in the plant vasculature contribute to maintaining the shoot K^+^/Na^+^ content ratio at a high value, which is a key determinant of salt tolerance (Munns and Tester, 2008; Maathuis, Ahmad and Patishtan, 2014; Ismail and Horie, 2017; Wu, Zhang, Giraldo and Shabala, 2018). In rice, as well as in Arabidopsis and various other species, clear evidence has been obtained that Na^+^ transporters belonging to the HKT family are involved in desalinization of the ascending xylem sap (Hauser & Horie, 2010). The H^+^/Na^+^ antiport system SOS1 has been suggested to also contribute to this function when the concentration of Na^+^ in the xylem sap reaches very high values (Maathuis et al., 2014). Regarding the mechanisms controlling K^+^ translocation to shoots, outwardly rectifying channels belonging to the Shaker family, among which SKOR in Arabidopsis and OsK5.2 in rice, have been shown to mediate K^+^ secretion into the xylem sap under normal conditions (Gaymard et al., 1998; Nguyen et al., 2017) but their contribution to this function under salt stress remains poorly documented. It should be noted that K^+^ secretion into the xylem sap may have to be active, under some environmental conditions (Wu, Zhang, Giraldo and Shabala, 2018), which would exclude channel-mediated (passive) contribution to this function in such conditions. Active H^+^-coupled K^+^ transports mediated by HAK/KUP/KT transporters (see below) or involving a NRT1/PTR family member, NRT1;5 (Li et al., 2017), have been shown to contribute to K^+^ translocation towards the shoots. In rice, OsHAK1 and OsHAK5, which are thought to be endowed with H^+^-K^+^ symport activity (Véry et al., 2014), have been shown to play a role in K^+^ translocation towards the shoots under saline conditions (Chen et al., 2015; Yang et al., 2014). The mechanisms that underlie these contributions remain however to be specified. H^+^-K^+^ symport activity in parenchyma cells bordering the xylem vessels would result in K^+^ retrieval from the xylem sap since the pH gradient between the sap and the cytoplasm is inwardly directed and thus favors K^+^ influx into the cells. It has thus been hypothesized that such H^+^-K^+^ symporters may allow K^+^ acquisition within the stele by parenchyma cells, and that this would result in a higher concentration of K^+^ in xylem-adjacent cells, and thus in an outwardly-directed K^+^ electrochemical gradient that would allow SKOR-like K^+^ channels to release K^+^ into the sap (Yang et al., 2014). At the leaf surface, control of stomatal aperture provides another type of contribution to salt tolerance. Exposure to saline conditions rapidly results in stomatal closure, which limits the flux of xylem sap, and thus the rate of Na^+^ translocation to shoots (Fricke et al., 2006; Huang et al., 2009; Hedrich and Shabala, 2018). Such a control is however likely to also affect the rate of K^+^ translocation to shoots, and thus its contribution to shoot K^+^/Na^+^ homeostasis should benefit from mechanisms allowing to counteract the depressive effect of the reduction in volumetric flow of xylem sap on K^+^ translocation.

OsK5.2, which belongs to Shaker channel subfamily 5 (outwardly rectifying Shaker channels) like its two counterparts in Arabidopsis SKOR and GORK (Véry et al., 2014), is expressed in both stomata and vascular tissues (Nguyen et al., 2017). Previous analyses have shown that OsK5.2 is involved both in xylem sap K^+^ loading, as SKOR in Arabidopsis (Gaymard et al., 1998), and in guard cell K^+^ release-mediated stomatal closure, as GORK (Hosy et al., 2003). The roles of SKOR and GORK in Arabidopsis salt tolerance remain poorly documented. The expression level of *OsK5.2* is fairly maintained in roots under saline conditions, and even increased in shoots in these conditions (Figure 2). This channel has thus been used as a model in the present report to investigate xylem sap K^+^ loading under salt stress, i.e., whether it can be channel mediated or requires active transport systems, and the involvement of stomatal aperture control in salt tolerance.

### 4.2 K^+^ Secretion into the xylem sap under salt stress

Saline conditions weakly affected the expression level of *OsK5.2* in roots (Figure 2a), as shown for its counterpart *SKOR* in Arabidopsis roots (Pilot et al., 2003). In line with this rather stable expression, the contribution of OsK5.2 to K^+^ secretion into the xylem sap (estimated from the difference in sap concentration between the WT and *osk5.2* mutant plants; Figure 4a) did not appear to be much modified by the salt treatment. This contribution even tended to slightly increase during the first week of the treatment, which may be due to increased driving force for K^+^ secretion under conditions of salt-induced membrane depolarization (Jayakannan et al., 2013; Mian et al., 2011).

Reliable measurements of both the membrane potential and the apoplastic K^+^ concentration of stelar cells are difficult to obtain. However, the fact that OsK5.2 can contribute to K^+^ secretion under salt stress provides definitive evidence that passive (since channel-mediated) secretion of K^+^ can occur in stelar cells of rice plants facing saline conditions. This conclusion, which does not exclude a contribution of active K^+^ transport mechanisms to K^+^ secretion under saline conditions, also means that other channels besides OsK5.2, either K^+^-selective and belonging to the Shaker family (Véry et al., 2014) or poorly K^+^-selective like NSCC channels identified in stelar cells by patch clamp experiments (Wegner & de Boer, 1997), could also contribute to K^+^ secretion into xylem sap under such conditions.

### 4.3 Reduction of the volumetric flux of xylem sap under salt stress

Exposure to saline conditions is known to rapidly result in reduced stomatal aperture and plant transpiration (Fricke et al., 2006; Hedrich & Shabala, 2018; Robinson, Véry, Sanders, & Mansfield, 1997). In the present study, the transpiration rate was similarly reduced in WT and *osk5.2* mutant plants at the end of the salt treatment, by about 50% under light conditions and 30% under dark conditions (Figure 3). The kinetics of the reduction in transpiration rate was however more rapid in WT than in *osk5.2* mutant plants.

Upon an increase in external medium salinity, abscisic acid (ABA) produced in response to the resulting osmotic stress is rapidly directed to guard cells, where it is expected to activate the PYR/PYL/RCAR-ABI1 PP2C phosphatase-OST1 SnRK kinase signaling pathway, leading to guard cell anion channel activation and stomatal closure (Hedrich & Shabala, 2018). The actual contribution of guard cell anion channels to the triggering of stomatal closure upon salt stress has however been little investigated so far. Likewise, the role in stomatal closure upon salt stress of the K^+^ outward channels acting as downstream effectors (Pandey et al., 2007; Schroeder et al., 2001) was still poorly documented. Indeed, although extensive analyses have concerned the integrated involvement of transport systems in regulation of guard cell turgor (Jezek & Blatt, 2017), little information is related to high salinity conditions (Lebaudy et al., 2008; Thiel & Blatt, 1991; Véry, Robinson, Mansfield, & Sanders, 1998). Here, our data reveal the important role of outward K^+^ channel activity in control of stomatal aperture upon salt stress. The whole plant transpiration data (Figure 3) indeed indicate that OsK5.2 activity in stomata contributed to the reduction in stomatal aperture observed over the entire duration of the salt treatment. Furthermore, our data indicate that this activity is of particular importance at the onset of salt stress by allowing a more rapid reduction of stomatal aperture (Figure 3).

### 4.4 Na^+^ and K^+^ translocation to shoots by the xylem sap and K^+^/Na^+^ shoot homeostasis

An apoplastic pathway strongly contributing to Na^+^ entry into the root and radial migration to the root vasculature (the so-called bypass flow across the root to the xylem) has been evidenced in rice in the presence of high Na^+^ concentrations (Faiyue, Al□Azzawi, & Flowers, 2012; Flam-Shephered et al., 2018; Yeo, 1998). In our experimental conditions, the concentration of Na^+^ in the xylem sap in both WT and *osk5.2* mutant plants was quite similar to that in the hydroponic medium (50 mM) (Figure 4). Also, the xylem sap K^+^/Na^+^ concentration ratio decreased very rapidly down to values lower than 0.1 after one day of salt treatment, and was then more than 10 times lower than the K^+^/Na^+^ root content ratio (Figure S2). This indicates that the former ratio (in the xylem sap) was not likely to reflect the corresponding ratio in the root symplasm. Altogether, these results support the hypothesis that the bypass flow of Na^+^ was the major determinant of the migration of this cation towards the xylem vasculature. In such conditions, the flux of Na^+^ translocated to the shoot becomes proportional to the volumetric flow of xylem sap. Since the Na^+^ concentration of the xylem sap was similar in the mutant and WT plants, the larger rate of transpiration under salt stress in the mutant plants due to impaired control of stomatal aperture (Figure 3) is the major determinant of the difference in Na^+^ translocation rate between the two types of plants (Figure S1b). Thus, OsK5.2-dependent control of stomatal aperture results in a reduction of Na^+^ translocation towards the shoots.

A reduction in the volumetric flow of xylem sap is however likely to impact the rate of K^+^ translocation to shoots (Figure S1a). Because OsK5.2 contributes to K^+^ secretion into the xylem sap, besides its involvement in stomatal aperture control, the reduction in the rate of K^+^ translocation to shoots induced by the salt treatment is more reduced in WT than in osk5.2 mutant plants. In other words, although OsK5.2 activity in stomata has a negative effect on K^+^ translocation to shoots by decreasing the transpiration rate (Figure 3), the contribution of OsK5.2 to K^+^ loading into the xylem sap (Figure 4a) outperforms the “negative” effect resulting from reduced xylem volumic flow (Figure S1a). This conclusion supports the hypothesis that the beneficial effect, in terms of control of Na^+^ translocation to shoots and tolerance to salinity, of the reduction in stomatal aperture upon salt stress is likely to integrate the plant ability to increase, or at least maintain, the rate of K^+^ secretion into the xylem sap.

Due to the beneficial effects of the overall OsK5.2 activity, the ratio of the K^+^ to Na^+^ translocation rates towards the shoots (identical to the xylem sap K^+^/Na^+^ concentration ratio; Figure 4c) is larger in WT than in *osk5.2* mutant plants. This is probably the main reason why the kinetics of the decrease in shoot K^+^/Na^+^ content ratio is slower in WT than in mutant plants (Figure 7b), and why the overall activity of OsK5.2 contributes to salt tolerance (Figure 1).

### 4.5 Contribution of a K^+^ channel to plant salt tolerance

Salt stress, not only strongly increasing shoot Na^+^ content, but generally also leads to severe K^+^ deficiency (Hauser & Horie, 2010; Marschner, 2011). Since insuring efficient root K^+^ uptake from soil appears as the primary way to insure shoot K^+^ feeding, most studies aiming at identifying salt tolerance determinants among K^+^ transport systems have focused on root uptake systems. Exposure to high salinity can substantially depolarize root periphery cells and make passive K^+^ uptake through inwardly rectifying K^+^ channels thermodynamically impossible (Rubio et al., 2020). High-affinity HAK/KUP/KT transporters, expected to rely on pH gradients created by the H^+^-ATPase pump to energize inward K^+^ fluxes through H^+^-K^+^ symport mechanism, are therefore considered as the main K^+^ transport systems taking part in root K^+^ uptake under high saline conditions. Several HAK transporters have been shown to be involved in root K^+^ uptake and thereby to contribute to plant salt tolerance: AtHAK5 in Arabidopsis (Nieves-Cordones et al., 2010), and OsHAK1, OsHAK5, OsHAK16 and OsHAK21 in rice (Chen et al., 2015; Feng et al., 2019; Shen et al., 2015; Yang et al., 2014). KO mutations in these different genes have been shown to result in reduced K^+^ uptake and root K^+^ content, and probably as a consequence, also in reduced K^+^ translocation to shoots and often reduced shoot K^+^ contents. Such defects could be observed upon salt stress but also in absence of saline treatment, and resulted in reduced plant growth in all conditions (Chen et al., 2015; Feng et al., 2019; Nieves-Cordones et al., 2010; Shen et al., 2015; Yang et al., 2014), Increased plant Na^+^ uptake was noted in some mutants (Shen et al., 2015), which could originate from higher root cell polarization (Nieves-Cordones et al., 2017). Also supporting the importance of HAK-mediated plant K^+^ uptake in salt tolerance, transcript level variations in the *OsHAK1* gene between rice subspecies have been found to underlie the difference in their salt tolerance (Chen et al., 2015).

It is also well known that transport systems from the H^+^/cation antiporter families, involved in K^+^ and Na^+^ intracellular compartmentalization, are major contributors to salt tolerance (van Zelm et al., 2020) through their roles in Na^+^ compartmentalization and turgor regulation, but also through indirect contributions to K^+^ homeostasis. For instance, increased activity (due to overexpression in transgenic plants) of the antiporter AtNHX1 from Arabidopsis or LeNHX2 from tomato has been shown to result in improved root K^+^ uptake and higher K^+^ contents in all tissues. Such effects, which are beneficial to salt tolerance, have been proposed to result from a decrease in cytosolic K^+^ concentration that these transport systems would generate, by compartmentalizing K^+^, which would lead to increased expression and/or activity of high affinity K^+^ transporters involved in root K^+^ uptake (Leidi et al., 2010; Huertas et al., 2013). Altogether, these studies provide evidence of strong interactions between K^+^ uptake, compartmentalization and translocation to shoots.

Other K^+^ transport-mediated mechanisms of plant salt tolerance and in particular mechanisms involving K^+^ channels, were reported but have not yet been deciphered. Transcriptional regulation of a few K^+^ channel genes, especially the strong up-regulation of the inward Shaker regulatory subunit *AtKC1* in leaves (Pilot et al., 2003), suggests a role of inward K^+^ channels in salt tolerance, which has not been determined so far. Here, we showed that KO mutation in the outward Shaker K^+^ channel gene *OsK5.2* leads to increased salt sensitivity. Lack of *OsK5.2* functional expression was found to result in impaired growth in plants subjected to saline conditions but not in plants grown in standard conditions (Figure 1), in contrast to what has been reported in KO mutant plants impaired in root K^+^ uptake, which mostly showed growth defects even in absence of salt stress (see above). OsK5.2 is involved in control of stomatal aperture and in K^+^ secretion into the xylem sap, and these two functions together underlie its contribution to salt tolerance. It is also worth to note that lack of OsK5.2 activity results also in impaired root K^+^ uptake under saline conditions since plant growth and root and shoot K^+^ contents were lower in *osk5.2* mutant plants compared with the corresponding WT plants (Figure 1 and Figure 5). Together with the defects in K^+^ translocation to shoots under salt stress that have been reported in mutant plants impaired in root K^+^ uptake or in K^+^ intracellular compartmentalization (see above), the reduction in K^+^ uptake resulting from lack of OsK5.2 channel activity provides evidence that the three functions, uptake, compartmentalization and translocation, are especially intensively coordinated under saline conditions. In conclusion, the present results highlight K^+^-channel-mediated mechanisms of salt tolerance, and provide a new possible target for plant breeders towards the improvement of tolerance to salt stress in rice.

## Acknowledgments

We are grateful to Emmanel Guiderdoni, Christian Chaine, Eve Lorenzini, Christophe Périn and Remy Michel for the rice mutant line amplification.

## AUTHOR CONTRIBUTIONS

A.-A.V., H.S., T.H.N. and J.Z. conceived the original research plans; A.-A.V., and D.T.L. supervised the experiments; T.H.N. and J.Z. performed the experiments. A.-A.V., H.S., T.H.N. and J.Z. analyzed the data; A.-A.V., H.S. and J.Z. wrote the first draft of the manuscript. All authors contributed to the article and approved the submitted version.

## SUPPORTING INFORMATION

Additional supporting information may be found online in the Supporting Information section at the end of this article.

**Table S1**: Primers used for qRT-PCR experiments

**Figure S1** K^+^ and Na^+^ fluxes arriving at light in leaves of wild-type and *osk5.2* mutant plants under control and salt treatment conditions.

**Figure S2** The ionic composition of the xylem sap does not reflect the K^+^ and Na^+^ relative contents of the roots.

