## Supplemental Figure S1 for "The outward Shaker channel OsK5.2 is beneficial to the plant salt tolerance through its role in K^+^ translocation and its control of leaf transpiration"

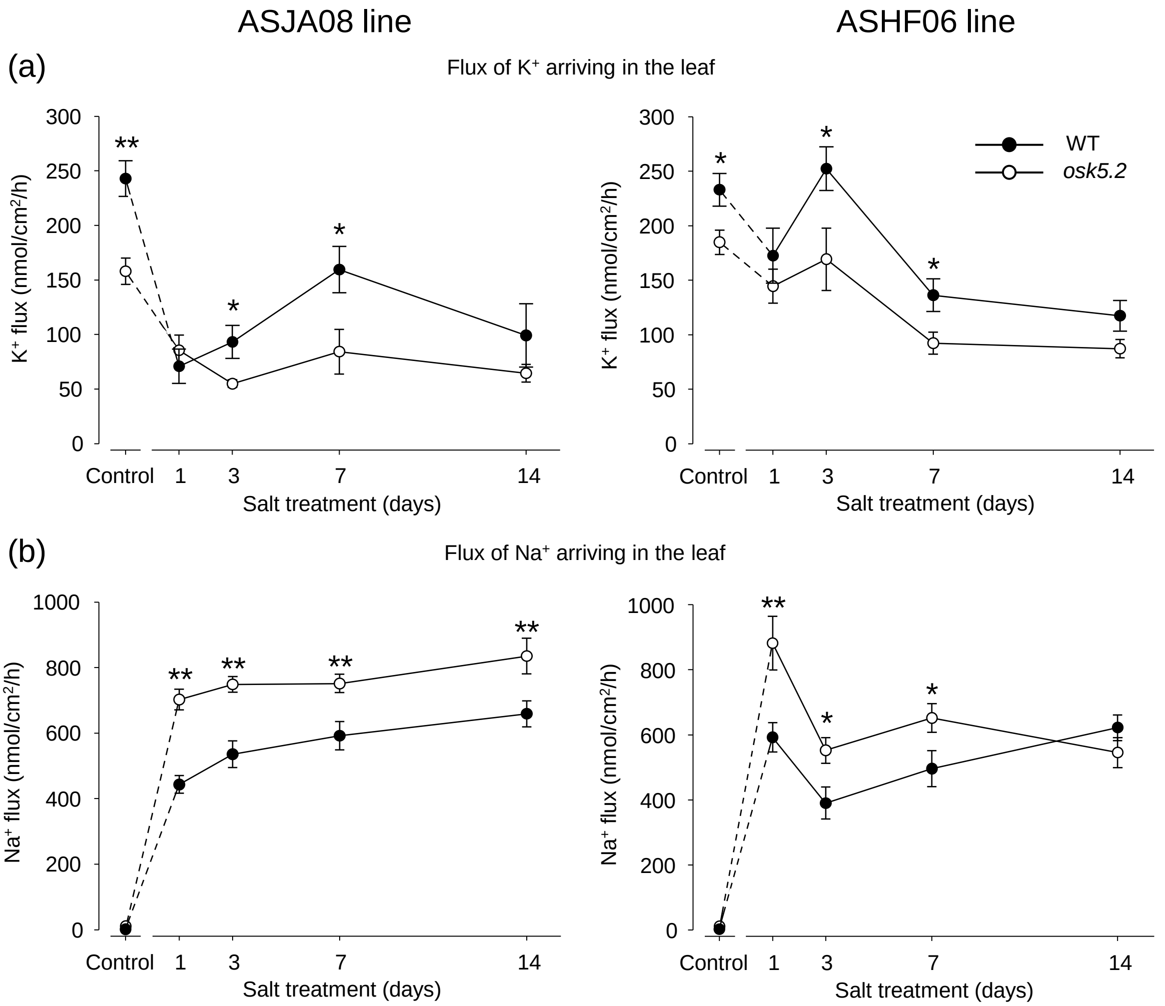


**Figure S1** $K^+$ and $Na^+$ fluxes arriving at light in leaves of wild-type and *osk5.2* mutant plants under control and salt treatment conditions. (a) and (b): normalized values (expressed per leaf surface) of $K^+$ (a) and $Na^+$ (b) fluxes assessed by multiplying the mean transpiration rates (data from Figure 3) by the corresponding $K^+$ or $Na^+$ concentrations in xylem sap (data of Figure 4). Left and right panels: *osk5.2* mutant plants (○) and corresponding wild-type plants (●) issued from ASJA08 (left) or ASHF06 (right) lines. Means ± SE; $n$ = 9 under salt treatment after 1, 3, 7 and 14 days, and $n$ = 12 under control conditions. See legends to Figures 3 and 4. Single and double stars denote statistically significant differences between the wild-type and *osk5.2* mutant plants (Student's $t$ test, $P \leq 0.05$ and $P \leq 0.01$, respectively).
