## Supplemental Figure S2 for "The outward Shaker channel OsK5.2 is beneficial to the plant salt tolerance through its role in K^+^ translocation and its control of leaf transpiration"

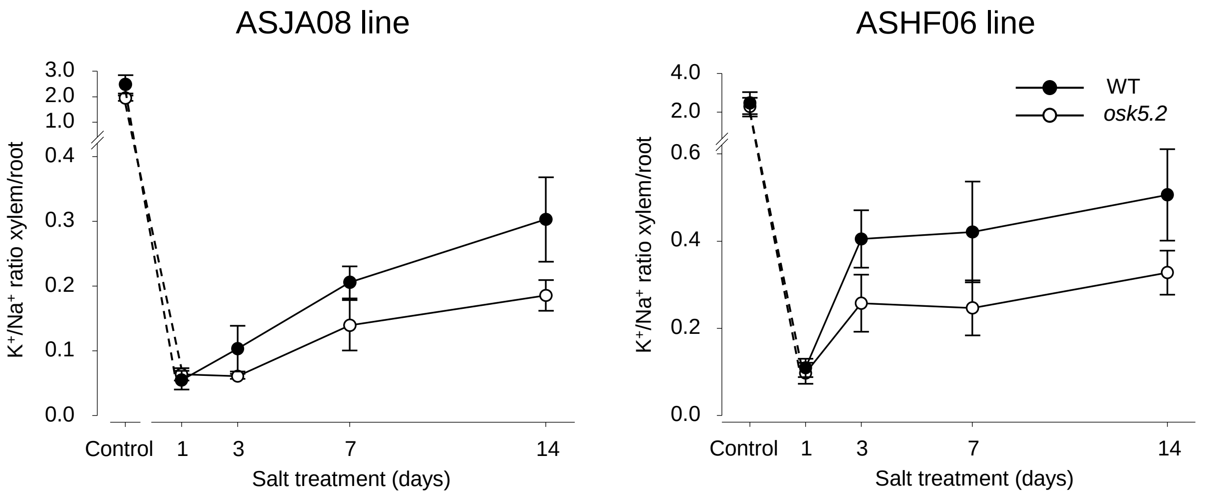


**Figure S2** The ionic composition of the xylem sap does not reflect the $K^+$ and $Na^+$ relative contents of the roots. Plants were subjected to a saline treatments (50 mM NaCl for 2 weeks; x axis). Y axis: xylem sap $K^+/Na^+$ concentration ratio (data from Figure 4c) reported to the root $K^+/Na^+$ content ratio (data from Figure 7a). Left and right panels: *osk5.2* mutant plants (○) and corresponding wild-type plants (●) issued from ASJA08 (left) or ASHF06 (right) lines. Means ± SE; n = 9 under salt treatment after 1, 3, 7 and 14 days, and n = 12 under control conditions. See legends to Figures 4 and 7.
