## Supplemental Table S1 for "The outward Shaker channel OsK5.2 is beneficial to the plant salt tolerance through its role in K^+^ translocation and its control of leaf transpiration"

**Table S1**: Primers used for qRT-PCR experiments.

| Gene | Primer name | Sequence (5'-3') | Amplicon length (bp) |
|---|---|---|---|
| *OsK5.2* | qPCR-OsK52-F358 | TTTGGCTTCTTCAGGGGGCT | 470 * |
| | qPCR-OsK52-R808 | TCTCCCAACTGCAAGCTCCC | |
| | qPCR-OsK52-F467 | CTGACACCTACCGCATGGTT | D-164 |
| | qPCR-OsK52-R611 | TCGTGTCAACCGAATCCACA | |
| *SMT3* | SMT3B303F | GGGAGGAGGACAAGAAGC | 273 * |
| | SMT3B303R | CTCCAGTCTGGTGGAGCAT | |
| | QRT94SMT3-F | CCTCAAGGTCAAGGGACAGG | D-94 |
| | QRT94SMT3-R | ACGGTCACAATAGGCGTTCA | |
| *Tip41* | Tip41B389F | TGGTTTTTGGGGAGAGTTTC | 389 * |
| | Tip41B389R | TGAAATGCCATTATCGGCTA | |
| | QRT101Tip41-F | TGGGAGTGATGCTTTGGTTC | D-101 |
| | QRT101Tip41-R | CAAGGTCAATCCGATCCTCA | |
| *eEF-1-beta2* | eEFB2Pcs | TGGTGAGGAGACTGAAGAGG | 418 * |
| | eEFB2Pcas | CTTCCGGATTTTTCTTTTTATC | |
| | qPCR-eEF1b2-F413 | TGGGAAGTCCTCAGTGTTGC | D-136 |
| | qPCR-eEF1b2-R549 | GTAACCAACCGGGACAAGCT | |

"D", pair of primers for transcript level quantification, that anneal on cDNA fragments derived from two exons. "*" refers to big amplicons that were used as a template for the amplification of calibration standards.
